# Neuromark dFNC Patterns: A fully automated pipeline to estimate subject-specific states from rs-fMRI data via constrained ICA of dFNC in +100k Subjects

**DOI:** 10.1101/2025.01.29.635539

**Authors:** M. Moein Esfahani, Vladislav Esaulov, Hemanth Venkateswara, Vince Calhoun

**Affiliations:** Department of Computer Science, Georgia State University, Atlanta, GA

**Keywords:** rs-fMRI, dFNC, constrained ICA, ICN, schizophrenia, spatial templates

## Abstract

Resting-state functional MRI (rs-fMRI) provides valuable insights into brain function during rest, but faces challenges in clinical applications due to individual differences in functional connectivity. While Independent Component Analysis (ICA) is commonly used, it struggles to balance individual variations with inter-subject information. To address this, constrained ICA (cICA) approaches have been developed using templates from multiple datasets to improve accuracy and comparability. In this study, we collected rs-fMRI data from 100,517 individuals across diverse datasets. Data were preprocessed through a standard fMRI pipeline. Our method first used replicable fMRI component templates as priors in constrained ICA (the NeuroMark pipeline), then estimated dynamic functional network connectivity (dFNC). Through clustering analysis, we generated replicable dFNC states, which were then used as priors in constrained ICA to automatically estimate subject-specific states from new subjects.This approach provides a robust framework for analyzing individual rs-fMRI data while maintaining consistency across large datasets, potentially advancing clinical applications of rs-fMRI.

## 1. INTRODUCTION

Today Resting-state functional MRI (rs-fMRI) has advanced our understanding of brain function in both typical and disordered states. By analyzing functional interactions using the BOLD signal, rs-fMRI offers key insights. Unlike task-based fMRI, which requires specific tasks, rs-fMRI captures spontaneous neural activity at rest[1, 2]. However, rs-fMRI faces significant challenges in clinical applications and single-subject analysis [1]. While task-based fMRI can rely on task-related features to guide analysis, rs-fMRI must identify intrinsic connectivity patterns that may vary considerably between individuals [3, 4, 5]. Researchers aim to understand brain organization through these interactions, but the true nature of functional patterns remains unclear. Unlike structural MRI, which relies on visible anatomy, rs-fMRI deals with hidden functional networks. A key challenge is identifying comparable functional patterns across individuals [1, 6]. In [1], Iraji et al. identified replicable networks to create intrinsic connectivity network (ICN) templates. We expanded this to dynamic functional network connectivity (dFNC) patterns, analyzing transient brain states. Our study calculated replicable dynamic states from over 100K subjects, aiding in individual dFNC pattern estimation. Using these templates, researchers can estimate and compare dFNC patterns across individuals, despite significant differences in functional connectivity [7]. Unlike existing methods, which struggle to balance individual variation with inter-subject consistency, our approach identifies general dFNC patterns. We demonstrated its effectiveness by using a CNN model to differentiate schizophrenia (SZ) from healthy control (HC) subjects based on estimated dFNC states.

### 1.1. Data driven approach

There are two main ways to analyze brain data [1, 8, 9]. The first estimates functional properties (e.g., brain connectivity) for individuals and compares them across samples. Early data-driven methods, like independent component analysis (ICA), follow this approach but may not always find the best functional patterns across individuals. The second approach, group-informed methods, uses group-level data to create templates, improving consistency and accuracy, especially for clinical use. A recent example is constrained ICA (cICA), which applies group templates to individual data for better comparability [1, 10]. This study applies cICA to dynamic connectivity in dFNC analysis.

## 2. MATERIALS AND METHODS

### 2.1. Dataset

We combined rs-fMRI data from more than 20 diverse public and private datasets, totaling 100,517 individuals [1]. These datasets varied in gender, age, handedness, and health conditions, and were collected using different scanners and imaging protocols. Our goal was to develop reliable templates from this large sample. While demographic influences will be explored in future work, we focused on quality control (QC) to ensure consistency without additional exclusions. The QC criteria, detailed in fig. 1B, included requirements for time points, head motion limits, and alignment to a standard template. We also checked spatial consistency using brain masks. Of the total sample, 57,709 individuals (57.4%) passed QC, while 42,808 (42.6%) were excluded. We generated templates from the QC-passed data in batches, confirming their reliability and reproducibility across batches.

**Fig. 1.**
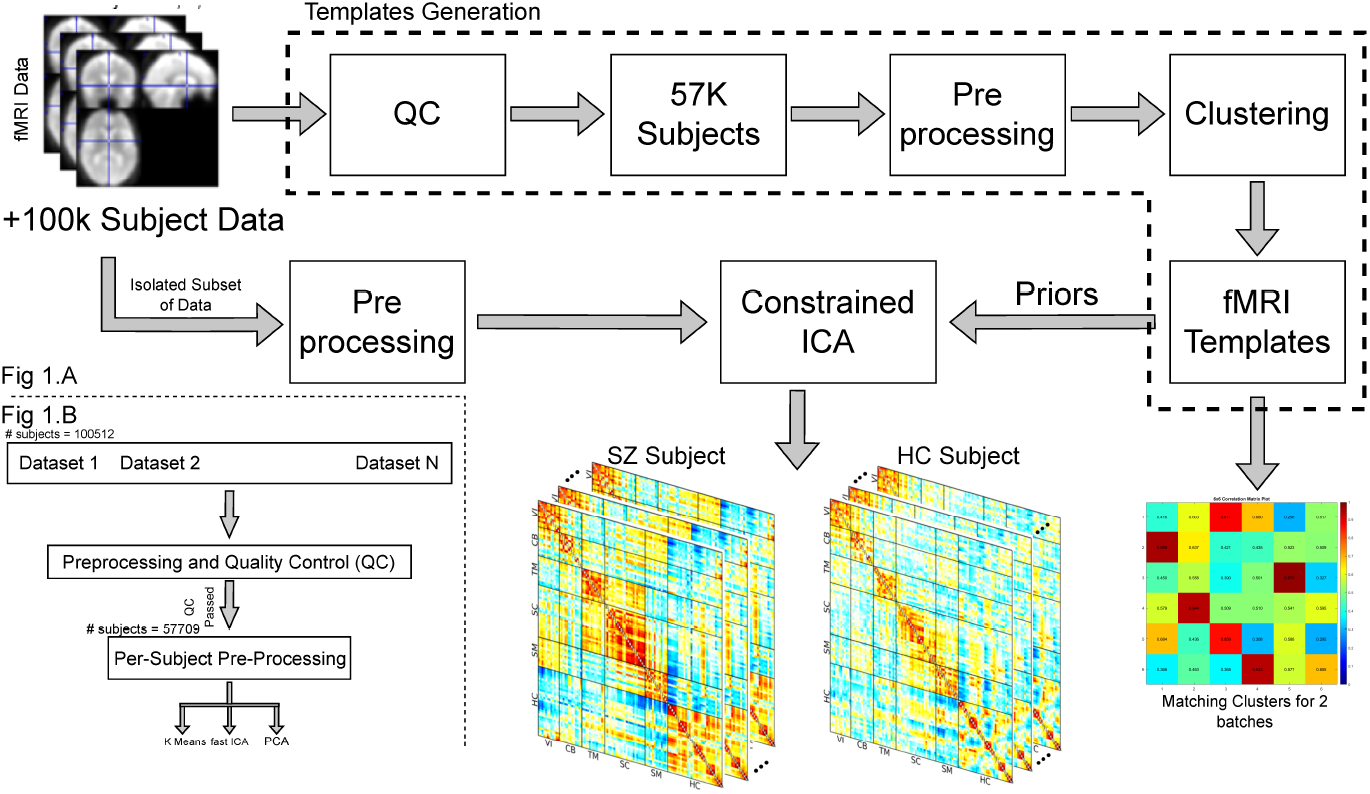
Schematic of the proposed pipeline, using extracted templates as priors for constrained ICA. The subsets of data were not part of QC and were isolated for testing, dataset has shuffled for multiple sites and information for generalization.

### 2.2. Preprocessing

To preprocess and clean the data, we used preprocessed data when it was available. If not, we applied our own steps based on fig. 1B. Figure 1B shows a diagram of this process. The first step was quality control, which removed subjects that didn’t meet the standards from [1]. After that, the second round of processing included steps like motion correction, slice timing correction, and distortion correction. For these tasks, we used tools from the FMRIB Software Library (FSL v6.0) and Statistical Parametric Mapping (SPM12) toolboxes within MATLAB. Distortion correction was done to fix warps caused by certain distortions using scans with different settings. This was done with FSL’s top-up tool using default settings. To handle noise and motion, we didn’t apply voxel-level regression-based correction. Instead, we relied on ICA to pick out and remove noise from the data. After preprocessing, we transformed the data into Montreal Neurological Institute (MNI) space using an echoplanar imaging (EPI) template[1, 11]. We chose the EPI template because it works better than structural ones, especially when distortion correction isn’t possible [1, 11]. We then resampled the data to 3 mm^3^ isotropic voxels and smoothed it with a Gaussian kernel of 6 mm full width at half maximum (FWHM). This preprocessing made sure the data was of high quality for further study. Each step, from motion and distortion correction to spatial normalization and smoothing, kept the data intact while reducing noise. A key point is that too much preprocessing could result in losing information, making the final templates less useful. This approach made sure the analysis was reliable, giving accurate and reproducible functional patterns.

### 2.3. Subject level preprocessing

The preprocessing method described above was applied to all subjects, regardless of the group or dataset. To further optimize results and ensure replicability, we tailored preprocessing for each subject based on their unique information. First, we applied high-pass and band-pass filters. Band-pass filtering (0.01 to 0.15 Hz) was done in two steps: a high-pass filter (using regressing cosines) followed by a low-pass filter with a Chebyshev Type II filter. To smooth the data and remove unwanted spikes, we used each subject’s data, including head motion, to interpolate and eliminate spikes using the General Linear Model (GLM). Since not all subjects experienced spikes or motion during recording, we applied this method across all subjects to ensure uniformity. We implemented the Chebyshev Type II filter for the high-pass step, setting the maximum loss in the passband to 3 dB and the minimum stopband attenuation to 30 dB. For the low-pass filter, we used a window size of 31. After filtering, we appended a DC set, which is a single cosine time series for frequencies between 0 and 0.01 Hz, defining the lower bound of the band-pass filter. For despiking, we used a method described on this page. This method adds noise-predictors to the design matrix, a technique used to model gradient artifacts that can cause spikes, a process known as despiking.

### 2.4. dFNC template estimation

Evaluating temporal coherence between intrinsic connectivity networks (ICNs) is widely used to assess both static and dynamic functional network connectivity (FNC) in the brain [3]. This method calculates functional connectivity (FC) at the network level and has shown promising results across multiple studies. In our study, FNC was calculated using data from 105 ICNs identified with the NeuroMark pipeline and the NeuroMark fMRI 2.1 template. Pearson correlation was used to analyze connections between ICNs, resulting in a 105×105 symmetric matrix with 5,460 unique values, providing a summary of static brain connectivity. Since dynamic FNC (dFNC) patterns vary within and across individuals, we used clustering methods like k-means, ICA, and PCA. Dimensionality reduction was applied to match the number of clusters, serving both as an analysis approach and a clustering method. In some cases, prior information can improve source estimation. Semi-blind ICA algorithms, like cICA, use this prior knowledge to refine results. The cICA algorithm [12] [13] incorporates constraints through a Lagrangian framework in the ICA objective function:

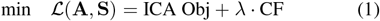

In our study, we identified replicable dFNC states across large datasets using ICA on windowed dFNC. These replicable states were then used as cICA constraints to automatically detect underlying states in new data while adapting to individual subjects. This approach improved both the estimation and consistency of the results across different datasets

## 3. RESULTS AND DISCUSSION

In this study, a sliding window approach was used for dFNC analysis to capture temporal changes in network connectivity patterns. A window length of 31 TRs with steps of 1 TR was chosen, shaped as a rectangular window of 31-time points convolved with a Gaussian kernel (sigma = 5 TRs) to create a tapering effect at the edges. This method enhanced the dynamic computation of dFNC from ICA time courses. To validate the dynamic analysis, all eigenvalues of the dynamic covariance matrices were checked to ensure they were positive, confirming stable and reliable covariance estimates.

The dFNC values were then Fisher-Z transformed to normalize their distribution and magnification was applied during visualization for better clarity. This method ensures accurate and meaningful connectivity patterns, accounting for individual and demographic differences to improve robustness and generalizability. To visualize brain state changes, the subject state transition vector (as shown in fig. 2) was used. This is a common concept in dFNC studies, tracking how brain connectivity shifts between cognitive or behavioral states over time. Instead of clustering all dFNC windows at once, clustering was first done on a subset of windows, known as subject exemplars, which showed maximal variability in correlations between component pairs. Exemplars were selected by identifying local maxima in variance across windows. On average, four to thirteen exemplars were selected per subject. The elbow criterion was applied to determine the optimal number of clusters, resulting in six centroid states (as depicted in fig. 3), within a search range of two to nine clusters. Correlation distance was used as the metric due to its sensitivity to connectivity patterns. In fig. 5, we visualized the templates that obtained as output of the results. These centroids served as starting points for clustering all dFNC windows, focusing on windows with the highest variability to ensure representative results. In fig. 4, we visualized the correlation of two batches (upper triangle 5460 points) obtained as output of K means clustering. This improved the robustness and accuracy of the clustering and can perform for better understanding of dynamic brain connectivity patterns.

**Fig. 2.**
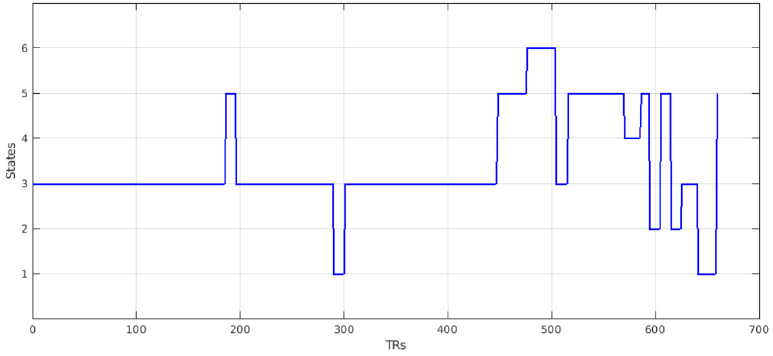
Subject state transition vector of a subject across a batch of subjects.

**Fig. 3.**
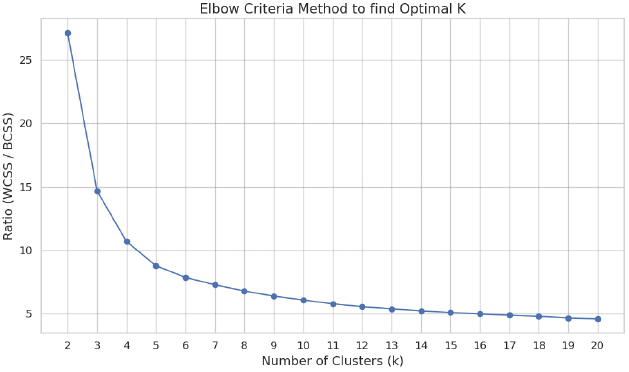
Elbow criteria to get the optimal number of states with K-means clustering.

**Fig. 4.**
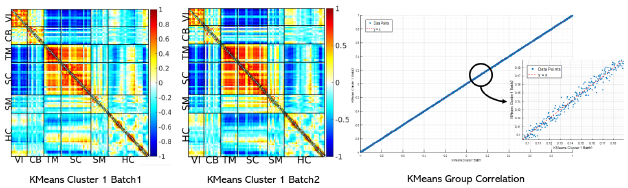
Two samples of data were selected, and group differences were depicted.

**Fig. 5.**
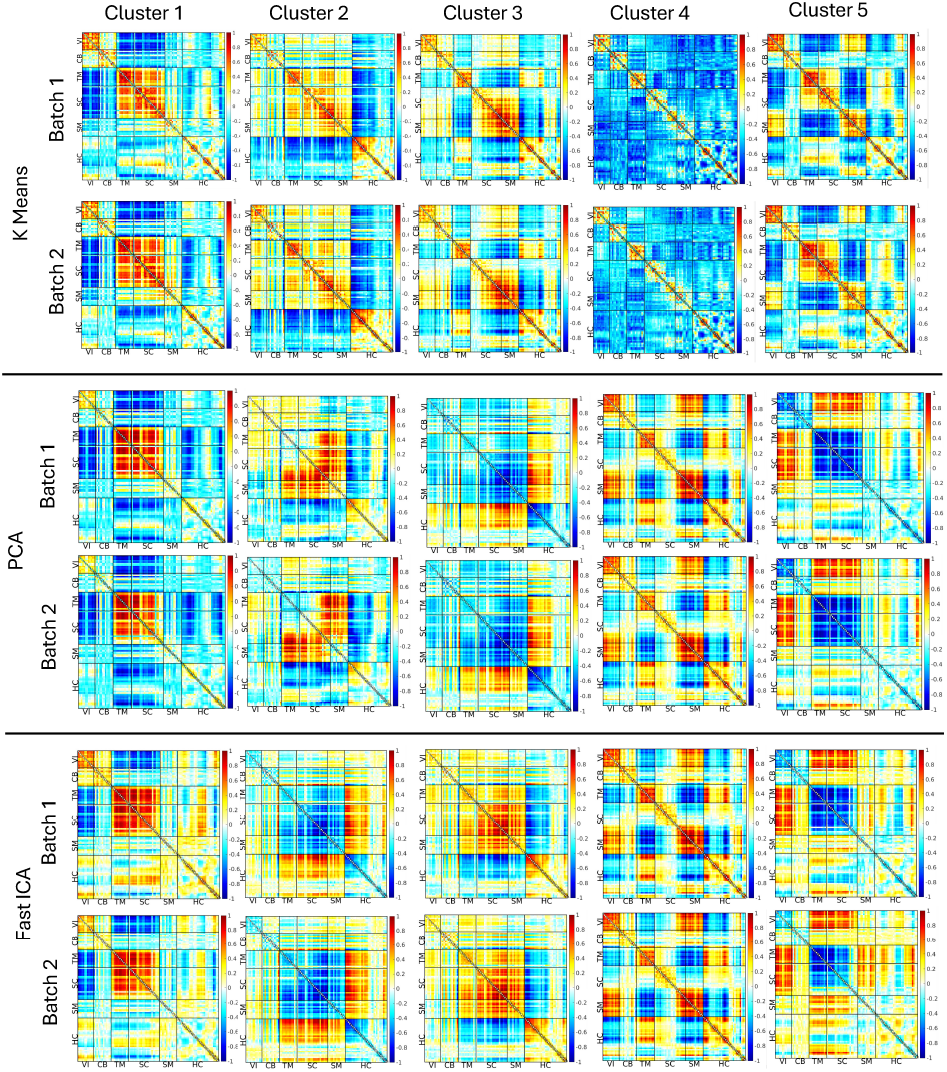
Results for templates obtained from K-Means, PCA, and fast ICA. ICNs are categorized into the following subcategories: visual (VI), cerebellar (CB), temporal(TM), subcortical(SC), somatomotor (SM), and Hippocampal (HC) networks. Every row represents a batch of results.

## 4. CONCLUSION

This study developed templates across multiple datasets to improve the accuracy and efficiency of analyses. By applying spatial cICA followed by dynamic cICA and incorporating prior knowledge from various datasets, more refined and consistent source estimations were achieved [4, 6, 14]. The findings demonstrate the value of using prior information in ICA across different research contexts. Future work will focus on studying the link between demographic variables and these findings, refining preprocessing methods, and enhancing the robustness and applicability of the results. Further research will explore dynamic connectivity patterns in neurological and psychiatric conditions, aiming for more effective interventions. Additionally, semi-blind ICA algorithms will be further examined to expand their generalization. In conclusion, ongoing efforts will refine these methods, advancing the understanding of brain connectivity dynamics and their implications for neurological and psychiatric conditions.

## 5. ACKNOWLEDGMENTS

This work was supported by grants from the NIH (R01MH123610) and NSF (2316421)

